# A waveform-independent measure of recurrent neural activity

**DOI:** 10.1101/2021.09.21.461229

**Authors:** Immo Weber, Carina R. Oehrn

## Abstract

Rhythmic neural activity, so called oscillations, play a key role for neural information transmission, processing and storage. Neural oscillations in distinct frequency bands are central to physiological brain function and alterations thereof have been associated with several neurological and psychiatric disorders. The most common methods to analyse neural oscillations, e.g. short-term Fourier transform or wavelet analysis, assume that measured neural activity is composed of a series of symmetric prototypical waveforms, e.g. sinusoids. However, usually the models generating the signal, including waveform shapes of experimentally measured neural activity are unknown. Decomposing asymmetric waveforms of nonlinear origin using these classic methods may result in spurious harmonics visible in the estimated frequency spectra. Here, we introduce a new method for capturing rhythmic brain activity based on recurrences of similar states in phase-space. This method allows for a time-resolved estimation of amplitude fluctuations of recurrent activity irrespective of or specific to waveform-shapes. The algorithm is derived from the well-established field of recurrence analysis, which has rarely been adopted in neuroscience. In this paper, we show its advantages and limitations in comparison to short-time Fourier transform and wavelet convolution using periodic signals of different waveform shapes. Further, we demonstrate its application using experimental data, i.e. intracranial electrophysiological recordings from the human motor cortex of one epilepsy patient.

## Introduction

During the last two decades, neural oscillations have gained increasing attention as a fundamental mechanism of neural communication (Buehlmann and Deco 2010; Agostina Palmigiano et al. 2017). Neural oscillations are defined as temporally recurring patterns of neuronal activity, also referred to as periodic and rhythmic activity. Oscillations mainly represent synchronized input to neural ensembles consisting of thousands of cells (Buzsáki and Draguhn 2004). The spatial specificity of recorded activity mainly depends on the measurement device used, with e.g. surface electrocorticography covering a much broader scale than e.g. invasive local field potential recordings (LFPs). Classically, oscillatory activity in the human brain is subdivided into five frequency bands: delta (<4 Hz), theta (4-8 Hz), alpha (8-13 Hz), beta (13-30 Hz) and gamma (>30 Hz) (Buzsáki and Draguhn 2004). A wide range of physiological processes in the animal and human brain is associated with fluctuations of oscillations in distinct frequency bands, e.g. such as deep sleep (delta oscillations, Amzica and Steriade 1998), long-term memory and inhibitory top-down control (theta oscillations, Oehrn 2018), attention and local inhibition (alpha oscillations, Bollimunta et al. 2011) and motor control (beta oscillations, Engel and Fries 2010). Further, alterations in distinct frequency bands occur in neurological and psychiatric diseases, e.g. changes in beta band activity during Parkinson’s disease (Hemptinne et al. 2013; Kühn et al. 2006; Little and Brown 2014) and theta activity during essential tremor (Pedrosa et al. 2012; Schnitzler et al. 2009). Nowadays, neuroscientists commonly use wavelet analysis for the quantification of oscillatory activity (Hramov et al. 2015; van Vugt et al. 2007) and Fourier-based analysis tools, such as multitapering or short time Fourier transformation (van Drongelen 2018). While being computational efficient, these methods have certain limitations that one needs to consider during interpretation. Classical Fourier and wavelet analysis usually implicitly assume that the analysed signal is a superposition, i.e. summation of stationary sinusoidal or wavelet shaped components. However, Fourier transformation of a non-periodic or non-sinusoidal signal may be difficult to interpret. While it is theoretically possible to deconstruct any non-periodic signal into a series of infinite sinusoidals using the Fourier transform, one has to be careful not to over interpret frequency components which arise due to the decomposition of a non-periodic non-sinusoidal signal into periodic sinusoidals (Lozano-Soldevilla et al. 2016; Gebber et al. 1999). Thus, the question arises which frequency components do indeed carry meaningful information, and which are redundant or even artificial.

The rationale behind using Fourier-based methods is the basic assumption, that most electrophysiologically recorded data, e.g. from electroencephalography (EEG) or LFPs, represent the summed activity of large neuronal populations (Franaszczuk and Blinowska 1985; Buzsáki and Draguhn 2004). However, despite few attempts at data-driven modelling of specific waveforms (Sherman et al. 2016; Lewis et al. 2012) most often the signal generating mechanisms and models and the prototypical waveform shapes of neuronal activity are unknown (Cole and Voytek 2017). In recent years, waveform shapes have gained increasing interest in the neuroscientific community (for reviews see Jones 2016 and Cole and Voytek 2017). Several studies revealed stereotypical variants of classic frequency bands which deviated from the sinusoidal waveform shape, e.g. the sensorimotor “mu rhythm” which is a variation of an alpha wave (Arroyo et al. 1993; Debnath et al. 2019; Muthukumaraswamy et al. 2004; Tiihonen et al. 1989) or motor cortical beta activity with a saw tooth shape (Cole et al. 2017). This non-sinusoidal rhythmic activity is functionally relevant. In Parkinson’s disease, asymmetric beta waves have been associated with the pathological state and shown to become more symmetric with successful treatment i.e. deep brain stimulation (Cole et al. 2017). While waveform shape is increasingly recognized to carry meaningful physiological information, there is still a lack of tools which specifically quantify non-sinusoidal activity. While recently, algorithms to characterize waveform shapes have been proposed, methods to incorporate non-sinusoidal activity into frequency analysis are still lacking (Cole et al. 2017; Pullon et al. 2019; Escobar Sanabria et al. 2017). Using classic approaches like Fourier analysis on asymmetric signals leads to the generation of harmonics in the respective spectra, which can be falsely interpreted as meaningful physiological or pathological activity. This is particularly true for measures of coupling, e.g. phase-amplitude coupling where non-sinusoidal signals may lead to spurious results (Lozano-Soldevilla et al. 2016; Yeh et al. 2016).

Here, we introduce a parsimonious way to analyse rhythmic activity that is not based on assumptions regarding waveform shapes. In this approach, we quantify recurrences of similar dynamic states ,based on the established framework of recurrence analysis (Webber and Marwan 2015). We derive a new algorithm for the estimation of a time-resolved recurrence amplitude spectrum and demonstrate its advantages and limitations in comparison to classic approaches, in particular in regard to non-sinusoidal signals. For this purpose, we use artificial data with known ground truth, as well as real intracranial brain recordings from the human motor cortex of one epilepsy patient.

## Methods

### 1. Basic definitions

For the remainder of this paper, let x_t_ be the realizations of stochastic variables X_t_ at time t generating a stochastic process *X*. Normal case letters indicate scalar valued observations, while bold letters indicate d-dimensional vector valued states. A state is defined as a collection of past mostly independent or temporally uncorrelated variables X_t-Δt,_ which are sufficient to predict the present observation X_t_. States can be reconstructed using Taken’s delay embedding theorem (Takens 1981) by time shifting the scalar time series *X* (d-1) times by a factor τ=Δt.

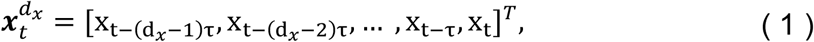

with *T* indicating the transpose of the vector. The dimension d represents the minimum number of degrees of freedom necessary to sufficiently describe the process *X*.

The collection of all realized states **x**_t_ is defined as the state-space of process *X*. For example, a perturbed frictionless pendulum creates a closed trajectory in a two-dimensional phase-space spanned by the variables position and velocity (Figure 1). The coloured dots in Figure 1B correspond to the pendulum positions shown in Figure 1A.

**Fig. 1:**
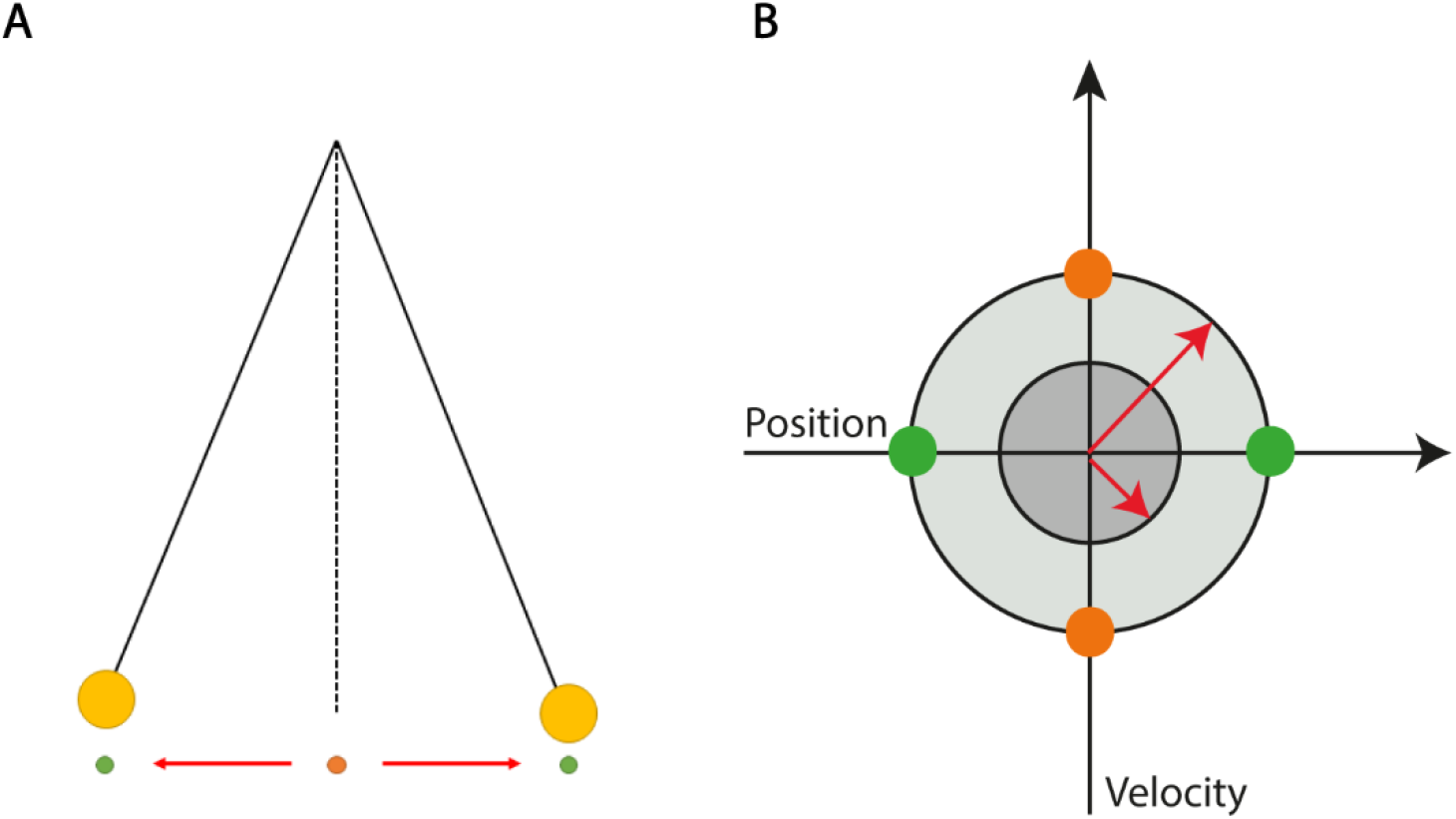
Example of a periodic phase-space representation. A: A frictionless pendulum perturbated from its resting position (orange) to a new position with small amplitude (green). B: Phase-space representation of the pendulum with two different amplitudes (grey-shaded circles). The state of the pendulum is uniquely determined by its momentary position and velocity. Colored circles in b correspond to the positions shown in a. Lengths of the red arrows indicate the absolute amplitudes for both phase-space representations.

Assuming the process *X* to be Markovian, i.e. stochastic with finite memory, the dimension d and the delay τ can be reconstructed from univariate time series using Ragwitz criterion (Ragwitz and Kantz 2002) or a combination of the false nearest neighbour algorithm (Hegger and Kantz 1999) and the auto-mutual information (Fraser and Swinney 1986) (for details see Supplement).

### 2. Quantification of recurrent states

#### 2.1 Frequency estimation: the recurrence period

A state **x**_t+Δt_ is defined to be recurrent after Δt time steps, if it is within a neighbourhood U_ε_ of X_t_ with radius ε (Eckmann et al. 1987):

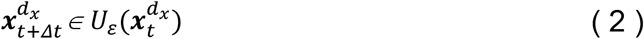

For infinitely small neighbourhoods, i.e. ε → 0, **x**_t+Δt_ is periodic with period Δt (Little et al. 2007). All recurrences of an arbitrarily high dimensional phase-space may be represented by calculating a two-dimensional binary recurrence matrix (Eckmann et al. 1987):

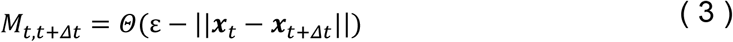

Θ is the Heaviside step function and ||.|| is the Euclidean distance norm:

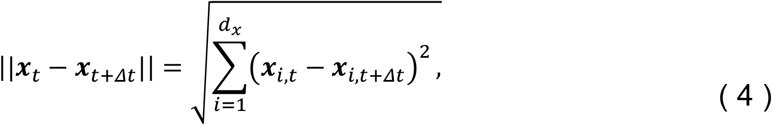

where i are the components of phase-space vectors. If (ε-||**x**_t_-**x**_t+Δt_||) is negative Θ is 0 else 1. This recurrence matrix M can be graphically represented by a recurrence plot, where each black dot represents one recurrence of time i at time j (Figure 2).

**Fig. 2:**
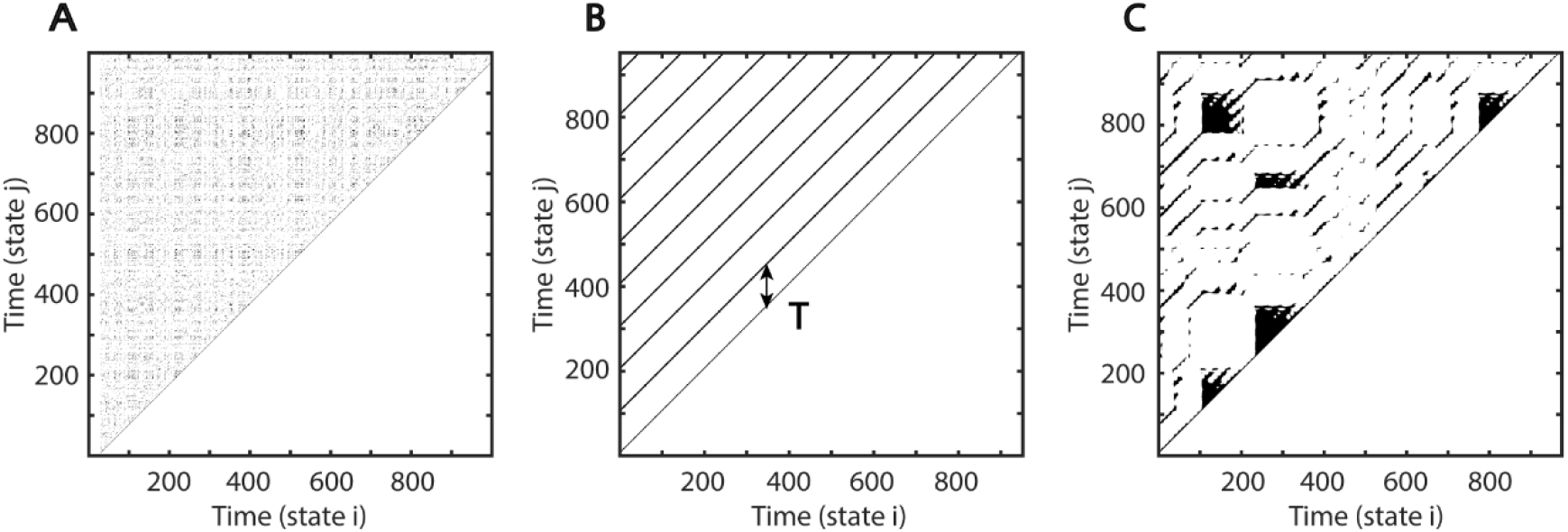
Examples of Recurrence Plots. At each time i each black dot indicates a spatial recurrence at time j. Parallel lines indicate periodic activity. Recurrence plots show characteristic patterns depending on the system’s qualitative dynamics: A: Random white noise. B: Sinusoid with period of 100 samples. C: Recurrence plot of a classic deterministic nonlinear system, i.e. the Lorenz system (Lorenz 1963).

Depending on the system’s local dynamics, the recurrence plot depicts different motifs. Parallel diagonal lines indicate periodicity and determinism while vertical lines appear due to laminar, i.e. unchanging behavior. White corners arise because of slow drifts or non-stationarity and isolated dots most often indicate stochastic behavior (Eckmann et al. 1987). The recurrence period T of any closest temporal neighbor x_t+Δt_ of x_t_ within a spatial neighbourhood U_ε_ may be estimated as the difference (Little et al. 2007):

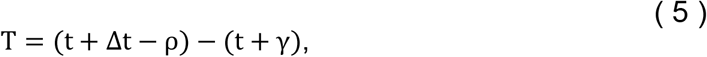

where γ is the difference in samples between **x**_t_ and **x**_t_ first leaving U_ε_ and ρ is the sample difference between x_t_ reentering U_ε_ and **x**_t+Δt_ (Figure 3A). In the recurrence matrix, T is equal to the number of states between vertical line segments starting from the main diagonal in M (see Figure 2B).

**Fig. 3:**
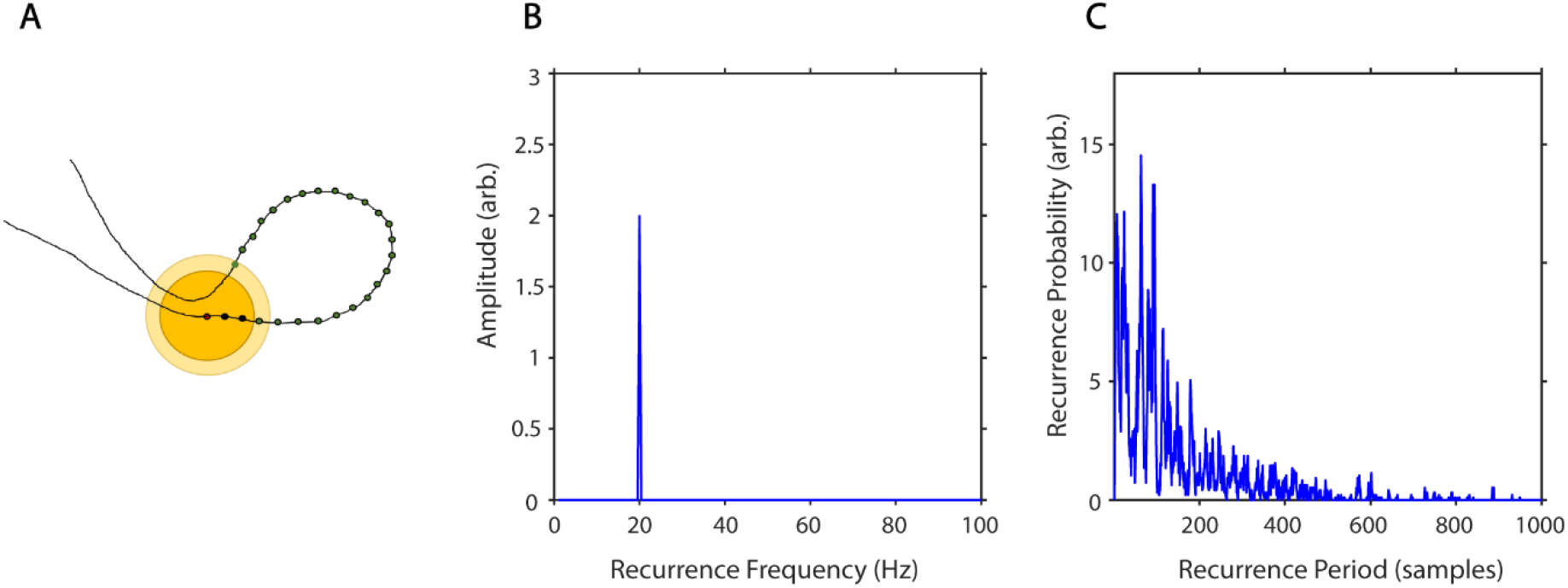
Estimation of recurrence periods. A: A closed local trajectory in phase-space. A reference point (red) is tracked in time until it reenters its spatial neighbourhood (examples of two neighbourhoods with different sizes in yellow). In this example the recurrence period is equal to 23, as it takes 23 samples (green) until the trajectory reenters the inner neighbourhood (dark yellow). B: Example estimation of recurrence amplitudes using the maximum norm. Recurrence amplitude of 5 s of a 20 Hz periodic signal with an amplitude of two. C: Example estimation of recurrence probabilities for a nonperiodic but highly recurrent deterministic system, i.e. the classic Lorenz system (Lorenz 1963). The plot shows the recurrence probabilities of each period T.

#### 2.2. Amplitude estimation

As can be seen in Figures 1B and 3A, recurrent systems form closed or nearly closed trajectories in phase-space. Here, the energy is contained in the phase-space volume of each recurrent state. In reference to our example, the phase-space portrait would increase in size, if the pendulum would be moved with a greater amplitude. Thus, a reasonable approximation of the amplitude of each period would be to estimate its maximum diameter in phase-space (see red arrows in Figure 1B).

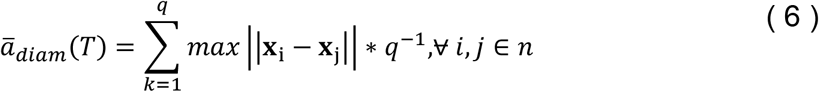

With q being the number of recurrences per T, i.e. how often a reccurence of duration T has occurred and n being the number of samples per recurrence (see Figure 3A).

This may be repeated for each recurrent trajectory per recurrence period T and subsequently averaged (Figure 3B).

#### 2.3. Recurrence probability

By estimating the recurrence period T for all phase-space vectors, it is possible to calculate a histogram R(T), where the bin number is equal to the longest recurrence period T_max_. The recurrence probability may thus be estimated by normalizing R(T) by the total number of recurrences (Figure 3C, Little et al. 2007). As noisy experimental data may lead to a high number of short period recurrences it is useful to calculate P(T) for a predefined range of T (T_min_ - T_max_):

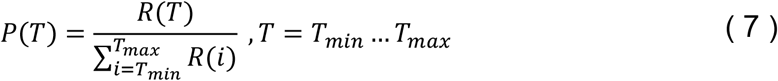

Finally, we weight each estimated mean amplitude by its recurrence probability to get an average for each recurrence frequency bin. This is equivalent to estimating the expected value of the amplitude for every recurrence frequency:

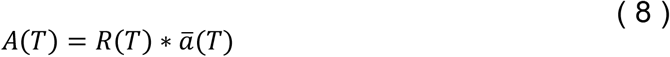

### Influence of neighbourhood ε on recurrence estimation

The recurrence probability is an estimate of the probability of a recurrence occurring after T time steps. The estimation of P(T) is dependent on ε. A choice of ε too small would result in many empty neighborhoods and thus in a high degree of statistical errors. If ε is chosen too large recurrences are not local anymore and recurrence periods are underestimated. The recurrence period gets approximately underestimated by two samples for every increase of ε by a multiple q of the sampling period fs (e.g. see the light yellow area in Figure 3A):

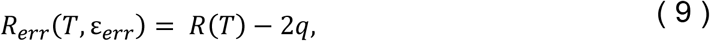

with

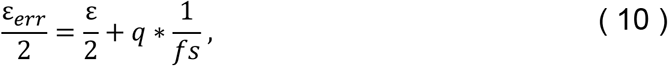

To better understand the influence of the parameter ε on recurrences, P(T) may be estimated over a wide range of neighbourhood-sizes, resulting in a spatially-resolved recurrence period spectrum (SREPS). In the SREPS, one would expect to find three regions of interest depending on ε. For very small ε the SREPS is governed by statistical errors and a uniform distribution across all T. For very high ε the distribution of P(T) is heavily shifted to small T with only few state vectors leaving and reentering any neighbourhood, with the extreme case of a neighbourhood-size fully engulfing the phase-space. If the signal contains any oscillatory activity slowly shifting but continuous “spectral” peaks occur in the intermediate range of ε. As stated above the best estimate of the recurrence period may thus be found at the crossing of the continuous spectral peaks and the noise regime for low ε (Figure 4).

**Fig. 4:**
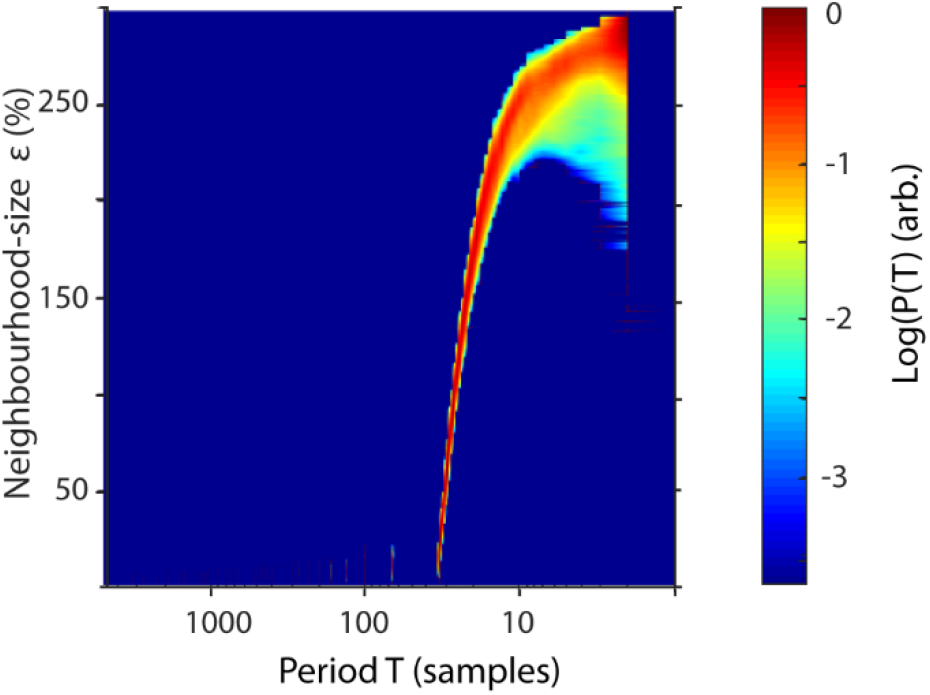
Spatially resolved recurrence probability. The plot shows recurrence probabilities P(T) of 50 s of a 3 Hz signal as a function of periods T=1-5000 samples and increasing neighbourhood-sizes ε=1-300. ε is represented in % of the standard deviation of the raw time series.

### 3. Time-resolved recurrence amplitude

In neuroscientific research it is often of interest to analyse spectral activity changes relative to some intervention, e.g. some stimulus or response. For this purpose, methods like short-term Fourier or Wavelet transform estimate power spectral density as a function of time (Hramov et al. 2015). Similarly, the recurrence amplitude spectrum may be estimated for n short overlapping time windows w_n_ of definite length:

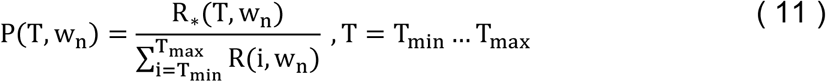

The length of each time window determines the maximum resolvable recurrence period and should thus be chosen with respect to the minimum frequency of interest.

### 4. Comparison between recurrence estimation, Fourier transform and wavelet transform

In Figure 5 we demonstrate the raw (Figure 5A) and weighted recurrence amplitude spectrum (Figure 5B) using an artificial signal of three signals of different concatenated waveform shapes, each with a length of five seconds, a frequency of 33 Hz and an amplitude of two: 1) a sinusoid, 2) a sawtooth wave and 3) a rectangle wave. While the frequency resolution succeedingly decreases for the three waveform types using the raw amplitude spectrum, it stays nearly the same for the weighted spectrum. However, the amplitude of the rectangle curve is slightly under- and its frequency slightly overestimated. Using the weighted amplitude spectrum increased the frequency resolution for the non-sinusoidal signals. For comparison with classic approaches, we analysed the same signal with short-time Fourier Transform (STFT, Figure 5C) and Wavelet analysis (Figure 5D). For the recurrence amplitude spectrum and the short-time Fourier spectrum we used 50 % overlapping windows of 600 ms lengths. For Wavelet analysis we used 30 cycles in order to approximate the window length for the other methods (1s). For STFT and Wavelet analysis, spurious harmonics can be seen for the non-sinusoidal waveform shapes in addition to the true frequency at 33 Hz. In contrast, the weighted recurrence amplitude spectrum shows only one spectral peak at a relatively narrow frequency band.

**Fig. 5:**
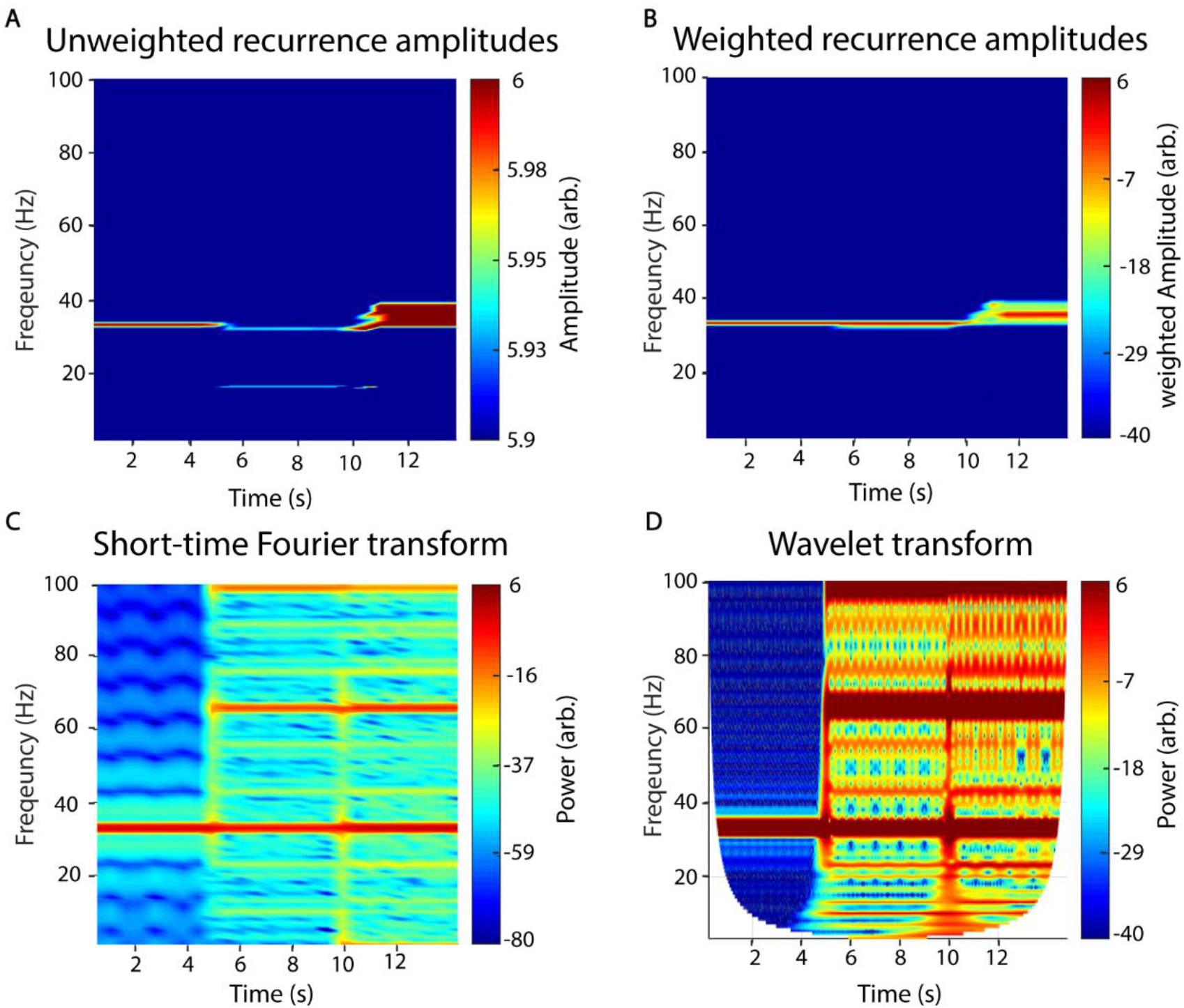
Recurrence amplitude spectrum. The recurrence amplitude spectrum of a compound signal of three 33 Hz oscillatory signals of different shapes is shown. The first five seconds are composed of a sine wave, the next five seconds of a sawtooth wave and the last five seconds of a rectangle wave. A: Raw unweighted amplitudes. B: Raw amplitudes were weighted with their respective probability densities. C: Short-time Fourier transform of the same signal. Window length was set to 1s with 50% overlap. Note the spurious harmonics for the nonlinear waveform shapes, i.e. rectangle (5-10s) and sawtooth shape (10 -15s). D: Wavelet transform of the signal. Wavelet width was set to 18. Note the spurious harmonics for the nonlinear waveform shapes, i.e. rectangle (5-10s) and sawtooth shape (10 -15s).

### 5. Waveform-specific filter

While the proposed estimator is independent of waveform, it is possible to tailor it to distinct waveform shapes. For this, we introduce a gain factor to the amplitude estimation step which is simply the Pearson correlation of the waveform shape of each recurrence and a scaled template waveform ζ.

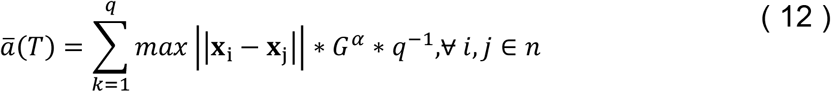

With α being chosen arbitrary and G being the gain factor:

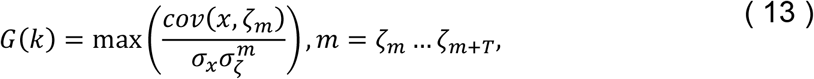

with cov being the covariance, σ being the variance and ζ_m_ being the template waveform from sample m to m+T. For perfectly matching waveforms, the gain is unity and thus the amplitude estimation is unaffected (Figure 6). In Figure 6 we used the same artificial signal as in Figure 5 and analysed it using waveform templates of a sinusoid (Figure 6A), sawtooth (Figure 6B) and rectangle shape (Figure 6C), respectively. The exponent α determines the strength of the gain factor and thus how much specific waveform shapes are filtered. In this example, it was set to five.

**Fig. 6:**
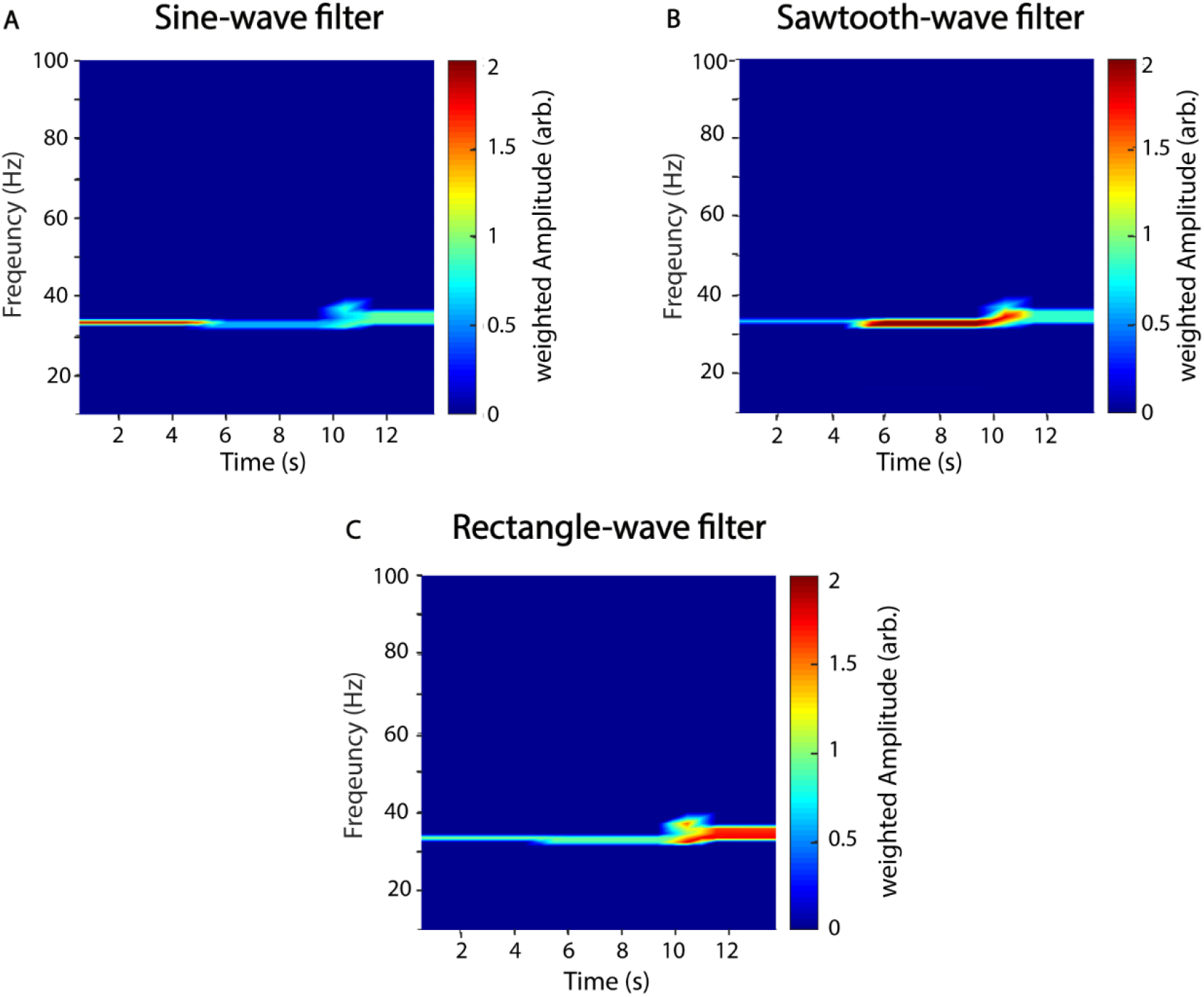
Recurrence amplitude spectrum with templates. The recurrence amplitude spectrum of a compound signal of three 33 Hz oscillatory signals of different shapes and a waveform dependent gain is shown. The first five seconds are composed of a sine wave, the next five seconds of a sawtooth wave and the last five seconds of a rectangle wave. Shown are the raw unweighted amplitudes. A: Five cycles of a sine wave were used as a shape template. B: Five cycles of a sawtooth wave were used as a shape template. C: Five cycles of a rectangle wave were used as a shape template. α was always set to 5. Window length was set to 1s with 50% overlap.

## Example application

While the recurrence amplitude spectrum might me estimated for all types of oscillatory activity, it is particularly suited for the analysis of electrophysiological neural activity. Thus, we briefly demonstrate its application using intracranial brain recordings, from the motor cortex of one epilepsy patient. The patient had received a 48 electrode electrocorticography (ECoG) grid for diagnostic purposes. Electrodes were localized by co-registration of pre-operative MRI and post-operative CT scans using the Fieldtrip toolbox (Oostenveld et al. 2011) (Figure 7A). As oscillatory activity in the motor cortex has been well characterized in previous studies, we selected three electrodes located on the precentral gyrus, i.e. the motor cortex (two medial electrodes in the proximity of the hand area and one lateral electrode). Anatomical selection was based on the AAL atlas implemented in FieldTrip (Tzourio-Mazoyer et al. 2002) (Figure 7A). We recorded electrophysiological data by means of the Neurofax-system of Nihon Kohden (Nihon Kohden, Rosbach, Germany) at a sampling rate of 1000 Hz. The patient participated in a study where he was asked to press a button on a standard computer keyboard. The study protocol was approved by the medical ethics committee Marburg and conducted in accordance with the latest version of the Declaration of Helsinki. To minimize edge effects occurring after convolution with a wavelet kernel, we segmented the data into relatively large time intervals of two seconds before until two seconds after response onset and discarded two seconds on each end of the segment after wavelet convolution. We removed line-noise, visually rejected artifacts and performed trend correction. We performed a response-locked data analysis, i.e. neural data is analysed time-locked to the patient’s button press. To this end, we used time-recurrence amplitude analysis and compared results to classic time-frequency analysis, i.e. wavelet convolution, in order to analyse power in different frequency bands.

**Fig. 7:**
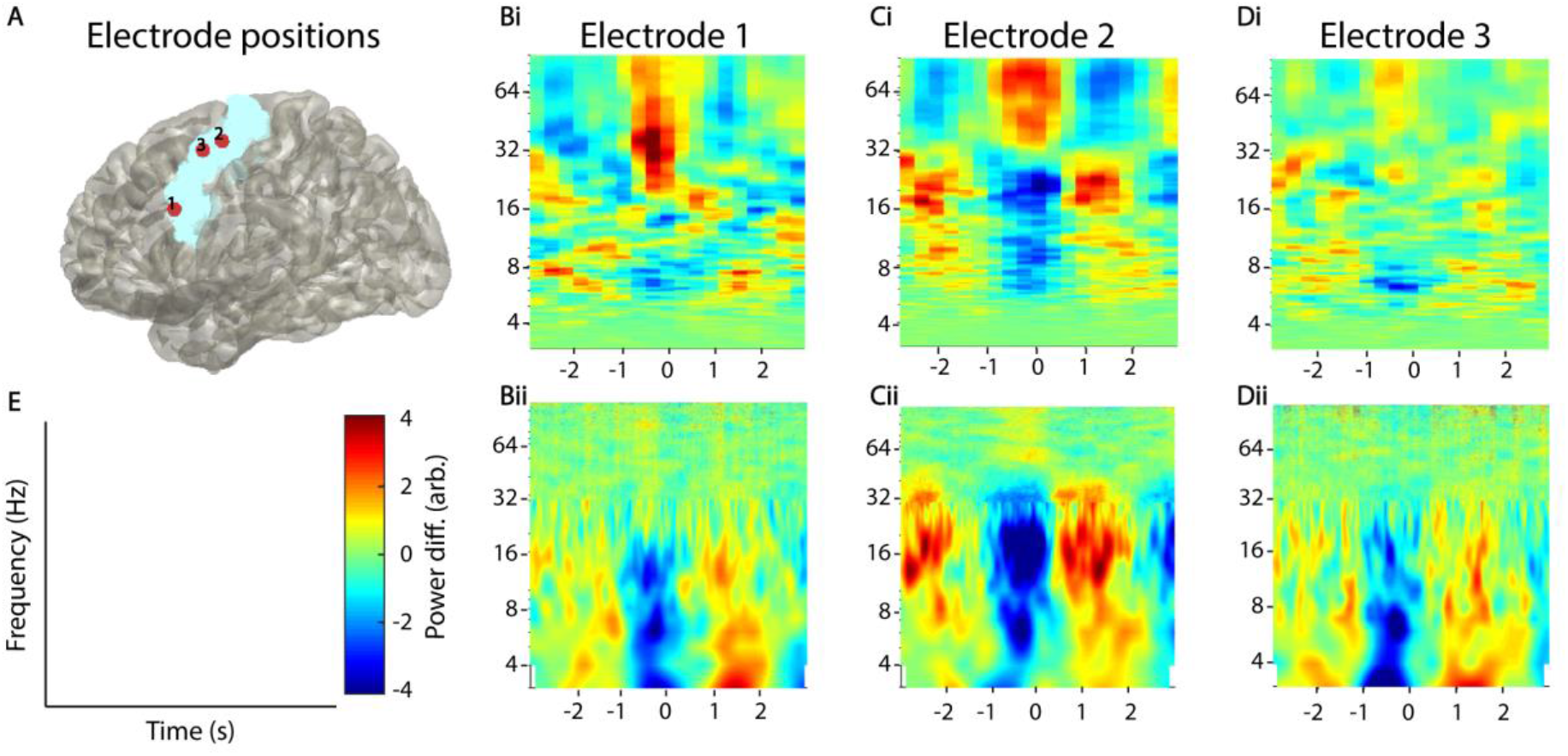
Example application on real invasive EEG data. A: Selected electrodes in the motor area of one epilepsy patient. The blue area indicates the precentral gyrus located with the AAL atlas in fieldtrip. Bi, Ci, Di: Recurrence analysis of three example electrodes. Plots have been convolved with a 5×5 smoothing kernel for visualization purposes. Bii, Cii, Dii: Corresponding Wavelet analysis of the same electrodes. All power values were normalized with respect to the total power of the analysed time window.

We used window lengths of 600 samples (=1.67s) and 50% overlap for the time-resolved estimation of recurrence amplitudes (Figure 7Bi, Ci, Di). Phase-space parameters, i.e. dimension d and delay τ were optimized for each time window using false nearest-neighbourhood algorithm and auto-mutual information, respectively. Neighbourhood-size was chosen ad-hoc at 70% of the standard deviation of each time series. For comparison with classic methods, we used a combined wavelet (for frequencies 3-30 Hz) and multitaper approach (for frequencies above 30 Hz; Figure 7Bii, Cii, Dii), as suggested for better frequency resolution and frequently adopted in the literature (van Vugt et al. 2007). To this end, we convolved the data with a continuous complex Morlet wavelet with seven cycles. From the wavelet-transformed signal, we extracted power values between 3 and 30 Hz in 1 Hz steps (lower frequencies are difficult to estimate using wavelet convolution in short time windows). For the multi-taper analysis, we used five tapers and a sliding 600 ms time window centered at one ms steps. Both spectra were normalized, using the complete time-period as baseline. Using wavelet analysis, we found a desynchronisation around the button press (∼-1 - 0 ms), ranging across frequency bands delta to beta (∼ 3-32 Hz) in all three electrodes (Figure 7CBii,Cii,Dii). After the motor response (∼ 1s post-response), we observed an enhancement in activity in this broad spectrum approximately one second after the response. This this so-called rebound activity occurred in a broad frequency range from delta to low gamma with it being most prominent in the beta band of electrode two. This beta rebound is a well-described phenomenon in the motor cortex during motor tasks (Crone et al. 1998; Pfurtscheller and Da Lopes Silva 1999; Jurkiewicz et al. 2006; Miller et al. 2007). Recurrence analysis showed a more specific beta desynchronisation and rebound for electrodes two, which was also more narrow band. In contrast to wavelet analysis, we additionally found a broad gamma activation during the button press, which was most prominently followed by a gamma desynchronisation in electrode two, but also visible in electrodes one and three to a lesser degree. In contrast, recurrence analysis did hardly reveal any changes in the theta band. Taken together, recurrence analysis was more sensitive for broad band high frequency activity that was not detected by wavelet convolution.

## Discussion

Here, we introduce a new method to analyse oscillatory activity in neural systems. We validated our approach with synthetic data and demonstrate its application with real experimental ECoG data of one epilepsy patient. Using artificial data, we show that the power estimates resulting from our method are more frequency-specific for non-sinusoidal waveforms and are not associated with spurious harmonics visible in the estimated frequency spectra. However, by applying specific waveform templates we demonstrate that our method is also able to detect specific shapes if necessary. Using human intracranial recordings from the motor cortex of one epilepsy patient, we show that recurrence analysis compares to conventional methods, such as wavelet analysis and Fourier transformation, in detecting movement-related oscillatory activity in the motor cortex.

With both methods, we found beta desynchronisation in the motor cortex around the button press followed by beta rebound (Salmelin et al. 1995; Jurkiewicz et al. 2006; Parkes et al. 2006). Both methods indicate that this effect is spatially specific, as it was most prominent in one of the two medial electrodes in proximity of the hand area of the primary motor cortex.

Further, recurrence analysis was more sensitive for detecting broad band gamma activity that is thought to represent multi-unit activity rather than narrow-band oscillations (Leszczyński et al. 2020). Recurrence estimation was seemingly less sensitive for theta and delta oscillations revealed by wavelet analysis. In contrast to beta activity in electrode two, theta/delta was more broadband in all three electrodes for wavelet analysis and much more narrowband for recurrence analysis. It is possible that this activity does not represent true oscillatory or recurrent activity, but rather local nonstationarities or drifts, which can hardly be detected with recurrence analysis. This is because states of local nonstationarities, even if they are oscillatory, do not come sufficiently close to each other after the full period to enter their respective neighbourhoods.

Though only meant as a proof-of-principle analysis, this is the first time directly comparing classic methods (i.e. wavelets and taper) with a recurrence based tool. Despite their fundamental differences in estimation procedures it is nevertheless evident that both capture recurrent neural activity, which validates both methodological concepts from different angles. However, discussing the mathematical equivalence of recurrence analysis and classic methods is beyond the scope of this study and should be analysed in future research.

Classically, neural oscillatory activity is analysed using methods derived from Fourier transform. Using these methods, time series get decomposed into prototypical waveforms or “wavelets” e.g. sinusoids. While this approach is justified by the understanding that e.g. EEG activity is a summation or superposition of thousands of synchronously active cells (Buzsáki and Draguhn 2004), interpretation of spectral estimation may be limited in some cases. This is because the generating models of neural activity and thus the basal waveform shape is most often unknown. Decomposing an asymmetric or nonlinear waveform into a series of sinusoids results in an infinite number of spurious harmonics (as demonstrated in Figure 5C-D), which may be misinterpreted as independent oscillatory activity. Thus, for such nonlinear signals classic techniques may generate a high degree of redundant information. Note, however, that wavelet analysis is in theory capable of redundandly quantifying nonsinusoidal oscillations by applying special mother wavelets e.g. like the Daubechies wavelets (Zhang et al. 2016). However, one drawback is that using a specific mother wavelet would still restrict analysis to one specific waveform shape and would also require prior knowledge, while recurrence based methods may detect unspecified arbitrary shapes. It is also still very uncommon to use any other mother wavelet than the Morlet wavelet for oscillatory analysis, although few studies exist that used other types to detect recurring spiking events in the EEG (Milton 1994; Grubov et al. 2017). The problem of nonsinusoidal waveform shape has been especially demonstrated for connectivity measures i.e. cross-frequency coupling (Yeh et al. 2016; Lozano-Soldevilla et al. 2016). Occurrences of nonlinear signals in electrophysiology are increasingly recognized to be commonly present in physiological (Arroyo et al. 1993; Gebber et al. 1999; Lozano-Soldevilla et al. 2016; Muthukumaraswamy et al. 2004) and pathological states (Cole et al. 2017). The physiological meaning of waveform shape, however, is still insufficiently understood (Cole and Voytek 2017).

The approach applied in this study utilizes the concept of recurrences in phase-space. The probability of a state recurrence as a function of its period has been previously introduced by Little et al. (2007). Our method builds upon this by also estimating amplitudes, i.e. energy content of a recurrence by estimating the phase-space volume of the recurrence. Additionally, by windowing the estimation procedure it is also possible to calculate a time-resolved recurrence amplitude spectrum, similar to short-time Fourier transform or wavelet analysis. The major difference to Fourier-based techniques is that time series are not fitted to basal waveforms in a “model-based” kind of way. Instead, it is quantified after what time the system reassumes a previous state, independent of the specific waveform in between these recurrences. Thus, as has been demonstrated in this study, simple asymmetric waveforms can be described more parsimoniously, without any spurious harmonics (Figure 5A-B). The most extreme example of this is a rhythmic idealized Dirac pulse, which has unity power over all frequencies in Fourier space (Beerends 2006), but can be parsimoniously represented with recurence based methods. Thus, another possible application of our proposed method might be analysis of spike train or electromyography data (EMG). The problems regarding the analysis of the latter with Fourier-based methods is widely recognized, which is why EMG data is often additionally preprocessed by e.g. extracting the Hilbert envelope or taking the absolute value (Myers et al. 2003). These preprocessing steps may, however, lead to spurious results depending on further analysis (Negro et al. 2015; McClelland et al. 2012).

### Parameters of recurrence analysis

While the concept of recurrences in phase-space is well established in other scientific fields e.g. climatology (Beaufort et al. 2001), proteomics (Webber et al. 2001) or geology (Donner et al. 2019), it is still scarcely applied in neurosciences. Examples include classification of mild cognitive impairment (Timothy et al. 2017), multiple sclerosis (Carrubba et al. 2019) or emotional states (Khodabakhshi and Saba 2020). One possible reason for this might be the number of parameters which need to be adjusted. While the algorithm is not parametric, i.e. not model-based *per se*, the estimation procedure may be sensitive to several key parameters due to the finiteness of measured data. These most prominently include the neighbourhood-size ε and the embedding parameters d and τ. For perfectly periodic recurrences and infinitely precise sampled data, the neighbourhood-size may be chosen arbitrarily small. However, for experimental data ε should be ideally chosen to barely engulf most of recurrent states. If the neighbourhood-size is chosen too large, recurrence periods get underestimated for every multiple of the sampling frequency. If ε is chosen slightly too small, recurrences might be missed. This may result in “harmonics”, i.e. multiples of recurrence periods appearing in the spectrum, as recurrences missed in one period might get detected in the next one (as can be seen in Figure 4). However, as our algorithm weights amplitudes by their probability of occurrences, few missed recurrences do not severely impact the overall spectrum. In the case that ε is chosen much too small, all meaningful recurrences might be missed and the spectrum gets dominated by measurement noise. The choice of ε thus depends on experimental data. However, it is important to note, that for real experimental data, there is no true neighbourhood-size as neural systems are hardly ever perfectly periodic. For intermediate ranges of ε recurrence spectra are rather stable and frequencies should only slowly shift (Figure 4). Nevertheless, when reporting results, neighbourhood and embedding parameters should always be reported for reproducibility. One possible approach to optimise neighbourhood-size is to estimate the recurrence amplitude spectrum as a function of ε and visually identify the noise regime for small neighbourhood-sizes (Figure 4). However, as this might be quite computationally demanding, it suffices to estimate the non-time resolved spectrum for a subsample of the data. This is justified if the variance of noise is static over time. The subset should be chosen long enough to cover the longest recurrence period of interest.

Of similar importance is an appropriate embedding of the measured data in phase-space. If the embedding dimension is too low, points which are far away from each other might get projected into close proximity. Thus, the time in between might be spuriously characterized as a specific recurrence period. On the other hand, if the embedding dimension is too high, estimation of recurrence periods becomes increasingly computational demanding and neighbouring points difficult to detect due to the increasing spaces between points otherwise known as “curse of dimensionality”. The embedding delay is important for spreading out the phase-space volume. A delay which is too small would result in all points laying on the first intersect and thus no closed trajectories to measure. For our algorithm we used the well-established false-nearest neighbours algorithm (Hegger and Kantz 1999) for the optimization of the embedding dimension and the auto-mutual information (Fraser and Swinney 1986). However, other techniques like e.g., the Ragwitz-algorithm are also frequently reported to optimize embedding parameters (Ragwitz and Kantz 2002; Michael Lindner et al. 2011; Weber et al. 2020, supplementary methods). By automatically optimizing d and τ we effectively eliminate these parameters, which makes the estimation procedure much easier to apply. This procedure is well established and implemented in many toolboxes utilizing the concept of phase-space analysis (Michael Lindner et al. 2011; Lizier 2014; Donges et al. 2015).

One limitation of the recurrence analysis is that it is by design not able to decompose a linear superposition of sine waves. For such artificially generated signals classic approaches are better suited. We thus propose to apply the demonstrated technique complementary in conjunction with classic approaches e.g. to discern possible spurious harmonics.

## Conclusion

In this study, we introduced a new time-resolved technique to measure amplitudes of oscillatory signals in a waveform independent manner. This method estimates the energy of recurrent activity by measuring distances of closed trajectories in phase-space, which are subsequently weighted by their respective probability densities. Using artificial data, we demonstrate that the measure generates less spurious harmonics due to nonlinear waveform shapes in comparison to classic techniques like Fourier Transform. Further, the analysis of intracranial data of one epilepsy patient indicates that recurrence might be better suited to estimate high frequency activity than congenital methods, such as wavelet analysis or the Fourier transformIn addition, we show that recurrence analysis can be used to specifically analyse signals with defined waveform shapes. In summary, the proposed measure might be well suited to complement classic frequency techniques, especially when the analysed signals are of nonlinear origin.

## Code availability statement

All code used in this study is readily available at https://nolitia.com/Code_recurrence_paper.zip.

