## Supplementary Methods for "A waveform-independent measure of recurrent neural activity"

#### Ragwitz criterion

Assuming the process *X* to be Markovian i.e. stochastic with finite memory, the dimension d and the delay τ can be reconstructed from univariate time series using Ragwitz criterion (Ragwitz and Kantz 2002). For a given combination of d and τ, neighbours of every state $x_t$ within a spherical neighbourhood $U_\varepsilon$ of diameter ε are iterated one time step. The mean of iterated neighbours is the prediction $\hat{\mathbf{x}}_{t+1}$ of $\mathbf{x}_{t+1}$.

$$\hat{\boldsymbol{x}}_{t+1}^{d_x} = \frac{1}{\left|U_\varepsilon(\boldsymbol{x}_t^{d_x})\right|} \sum_{\boldsymbol{x}_{t-\Delta t}^{d_x} \in \boldsymbol{U}_\varepsilon(\boldsymbol{x}_t^{d_x})} \boldsymbol{x}_{t-\Delta t+1}^{d_x} \quad \text{( S1 )}$$

Where t-Δt indicates that spatial neighbours temporaly precede $\mathbf{x}_t$ and |.| indicates the number of neighbours in $U_\varepsilon$. The combination of d and τ is chosen for which the root mean squared prediction error is minimum:

$$RMSPE = \sqrt{\frac{\sum_{t=1}^{n}\left(\hat{\boldsymbol{x}}_{t+\Delta t}^{d_x} - \boldsymbol{x}_{t+\Delta t}^{d_x}\right)^2}{n}} \quad \text{( S2 )}$$

where n is the number of states predicted.

#### False nearest neighbours algorithm

For deterministic systems, the phase-space dimension may be alternatively reconstructed using the false nearest neighbourhood method (FNN, Hegger and Kantz 1999):

$$\mathrm{FNN(d)} = \sum \frac{\theta\left(\frac{\left|\mathrm{x}_{\mathrm{i}+1} - \mathrm{n}(\mathrm{x}_\mathrm{i})_{\mathrm{j}+1}\right|}{\left\|\mathbf{x}_\mathrm{i}^\mathrm{d} - \mathbf{n}_{\mathrm{xi}}^\mathrm{d}\right\|} - \mathrm{Rtol}\right)}{\wedge\ \theta\left(\sqrt{\left(\left\|\mathbf{x}_\mathrm{i}^\mathrm{d} - \mathbf{n}_{\mathrm{xi}}^\mathrm{d}\right\|\right)^2 - \left(\left|\mathrm{x}_{\mathrm{i}+1} - \mathrm{n}(\mathrm{x}_\mathrm{i})_{\mathrm{j}+1}\right|\right)^2} - \mathrm{SD(X)} * \mathrm{Atol}\right)}, \quad \text{( S3 )}$$

with θ indicating the Heaviside-step function, i being the temporal index of x, j representing the temporal index of n, $n(x_i)$ indicating the next neighbor of $x_i$, ||..|| indicating the maximum norm, $R_{tol}$ being a distance threshold, $A_{tol}$ being a loneliness threshold and SD indicating the standard deviation. If the embedding dimension is too low, false neighbours of points in phase-space may arise due to projections. The optimal embedding dimension is thus the dimension for which the percentage of false neighbours drops to zero.

#### Auto-mutual information

The embedding delay may be estimated using the auto-mutual information (AMI, Fraser and Swinney 1986). Here, the shared information between the present and past of a process X is calculated as a function of a delay Δt. The Δt for which AMI drops to zero may be used as τ for the embedding:

$$AMI(\mathrm{X_t}; \mathrm{X_{t-\Delta t}}, \Delta \mathrm{t}) = \mathrm{H}(\mathrm{X_t}) - \mathrm{H}(\mathrm{X_t} | \mathrm{X_{t-\Delta t}}), \tag{S4}$$

with

$$\mathrm{H}(\mathrm{X}) = -\sum \mathrm{p}(\mathrm{X} = \mathrm{x}) \log_2 \mathrm{p}(\mathrm{X} = \mathrm{x}) \tag{S5}$$

and p(X=x) being the probability of X taking on the value of x.

### Publication bibliography

Fraser; Swinney (1986): Independent coordinates for strange attractors from mutual information. In *Physical review. A, General physics* 33 (2), pp. 1134–1140. DOI: 10.1103/physreva.33.1134.

Hegger, R.; Kantz, H. (1999): Improved false nearest neighbor method to detect determinism in time series data. In *Physical review. E, Statistical physics, plasmas, fluids, and related interdisciplinary topics* 60 (4 Pt B), pp. 4970–4973. DOI: 10.1103/physreve.60.4970.

Ragwitz, M.; Kantz, H. (2002): Markov models from data by simple nonlinear time series predictors in delay embedding spaces. In *PHYSICAL REVIEW E* 6505 (5), p. 6201.
